# Rakaia: interactive discovery of spatial biology at scale

**DOI:** 10.1101/2025.08.15.670550

**Authors:** Matthew Watson, Golnaz Abazari, Edward L.Y. Chen, Tiak Ju Tan, Jennifer L. Gorman, Caroline Diorio, Hal K. Berman, Kieran R. Campbell, Hartland W. Jackson

## Abstract

Spatial biology data throughput currently outpaces interpretable analysis, limiting large-scale discovery and translation. Here we present Rakaia, a browser-based platform for multiplexed imaging and spatial transcriptomics, empowering code-free interactive exploration and evaluation. Rakaia enables visualization, annotation, and feature-based querying with prioritization across thousands of images. We use Rakaia to identify cell types of interest in >200 highly multiplexed images from non-malignant human breast tissue. Rakaia is available at: https://rakaia.io/

## Main

Spatially resolved ‘omics technologies including antibody or probe-based multiplexed imaging [1, 2, 3], spatial transcriptomics (ST) [4, 5] and mass spectrometry imaging [6] generate comprehensive measurements as spots, beads, or pixels across single cells and tissues [7], while retaining spatial information. They have provided insights into cellular phenotypes within heterogeneous tissue architectures [8, 9] across a range of disease states and dynamic conditions [10, 11], enriching detailed visual discovery while complementing simple diagnostic standards such as hematoxylin & eosin (H&E) staining [12]. As a result, such assays are being increasingly adopted as core research components, capturing critical cellular and morphological information in studies spanning experimental conditions, patient samples, and even research sites [13, 14, 15, 16]. In response to broader community adoption of spatial research, consortia maintaining databases such as HuBMAP [17], CellxGene [18], and HTAN [19] are curating datasets on the order of thousands of images and spatial features totalling multiple terabytes in size, collating rich multimodal resources for generating novel hypotheses. Increased throughput combined with high data complexity has led to barriers for interpretability, reflecting the need for interactive user-interfaces (UIs) that can support user-guided visual comparisons to propose and confirm research aims. In particular, interactive visual inspection of many images, regions of interest (ROI), or fields of view (FOV) is a necessary step in spatial analysis workflows, enabling human-in-the-loop assessment to quantitatively compare and interpret differences in spatially defined features. As such, software that facilitates both visual exploration of large-scale datasets, as well as robust, code-free analysis to interrogate multiple data modalities, is fundamental for advancing integrated, standardized discovery beyond basic inspection of individual images.

Current graphical user interfaces (GUIs) for spatial analysis are limited to viewing one image at a time [20, 21] and lack features that enable interaction with multi-image datasets. Further, existing software can be limited to vendor-specific file types and platforms [22, 23] or may retain a high barrier for data import [24, 25, 26]. Image editing tools (e.g. gradient scaling, noise filtering, parameter standardization) and UI components to generate publication-quality images suitable for detailed interpretation are restricted [25, 27, 28], and fail to provide persistent storage of fine-tuned user selections [24, 27, 28]. Therefore, a single platform that accessibly unifies the necessary tools for both high-quality visualization and interpretation across hundreds to thousands of multiplexed spatial images is required.

To address this we developed Rakaia, a browser-based application optimized for analyzing spatial data at scale. Rakaia’s interactive UI enables users to visualize a variety of spatially-resolved technologies, implementing on-demand loading to enable rendering and comparisons of up to thousands of images in a single session. Rakaia’s integrated toolchain enables multi-image query/gallery display, object detection, quantification/clustering, and whole slide image (WSI) alignment (**Fig. 1**). Upon generating high-quality multiplexed images, users can layer matched cell/object segmentations, custom annotations, and perform image ranking to prioritize images for cohort-level spatial analysis (**Fig. 1**), leveraging a complete suite of tools to render, inspect, and analyze diverse spatial datasets.

**Figure 1.**
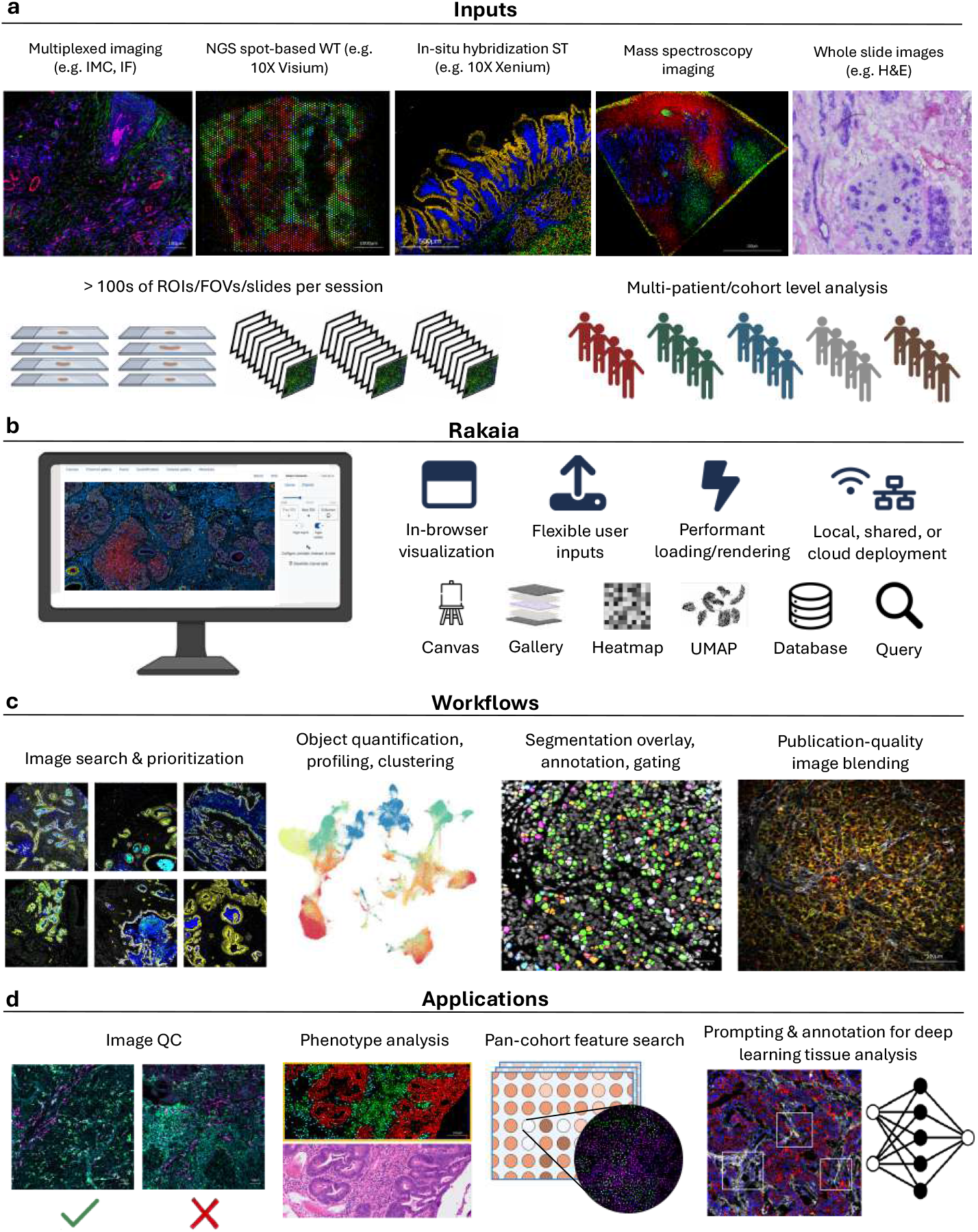
Overview of Rakaia. **(A)** Rakaia accepts a variety of common spatial assay types spanning proteomics, microscopy, ST assays and WSIs. **(B)** Rakaia’s in-browser GUI provides users with comprehensive tooling for complete analysis of multiplexed data **(C)** Unified workflows in Rakaia enable focused querying across multiple ROIs by leveraging object segmentation, quantification, and annotation. **(D)** Rakaia can be used to generate, validate, and communicate hypotheses for spatial datasets at scale, and outputs are extensible to broad sets of downstream tasks.

We present Rakaia as a user-accessible platform for researchers and clinicians to effectively discover and analyze spatial biology data, accommodating both routine image inspection as well as systematic and hypothesis-driven discovery and validation. In addition to enabling visualization scaled to biological variation, Rakaia facilitates the integration of both user annotation and quantitative analysis methods for effective comparisons of experimental conditions or identification of unique phenotypes within large sample-sets using the following built-in tools:

### Browse, search, and prioritization by image gallery

Rakaia includes a dataset gallery (**Extended Data Fig. 1**) where users can preview thumbnails from randomly assorted or ranked images. Ranking may be performed with phenotype-based searches to show the presence of similar structures or cellular phenotypes compared to a query image set or quantitative trait i.e. single cell cluster, thereby enabling prioritization of regions or cell types of interest (**Fig. 2**). Basic filtering functions alternatively allow users to search based on keyword patterns such as a list of patient identifiers or shared meta-features i.e. treatment or subgroup. The combination of gallery tab and querying functions enables users to browse and inspect thousands of images in a directed manner, with a single-click option to rapidly toggle to select images for full-focus visual inspection and annotation.

**Figure 2.**
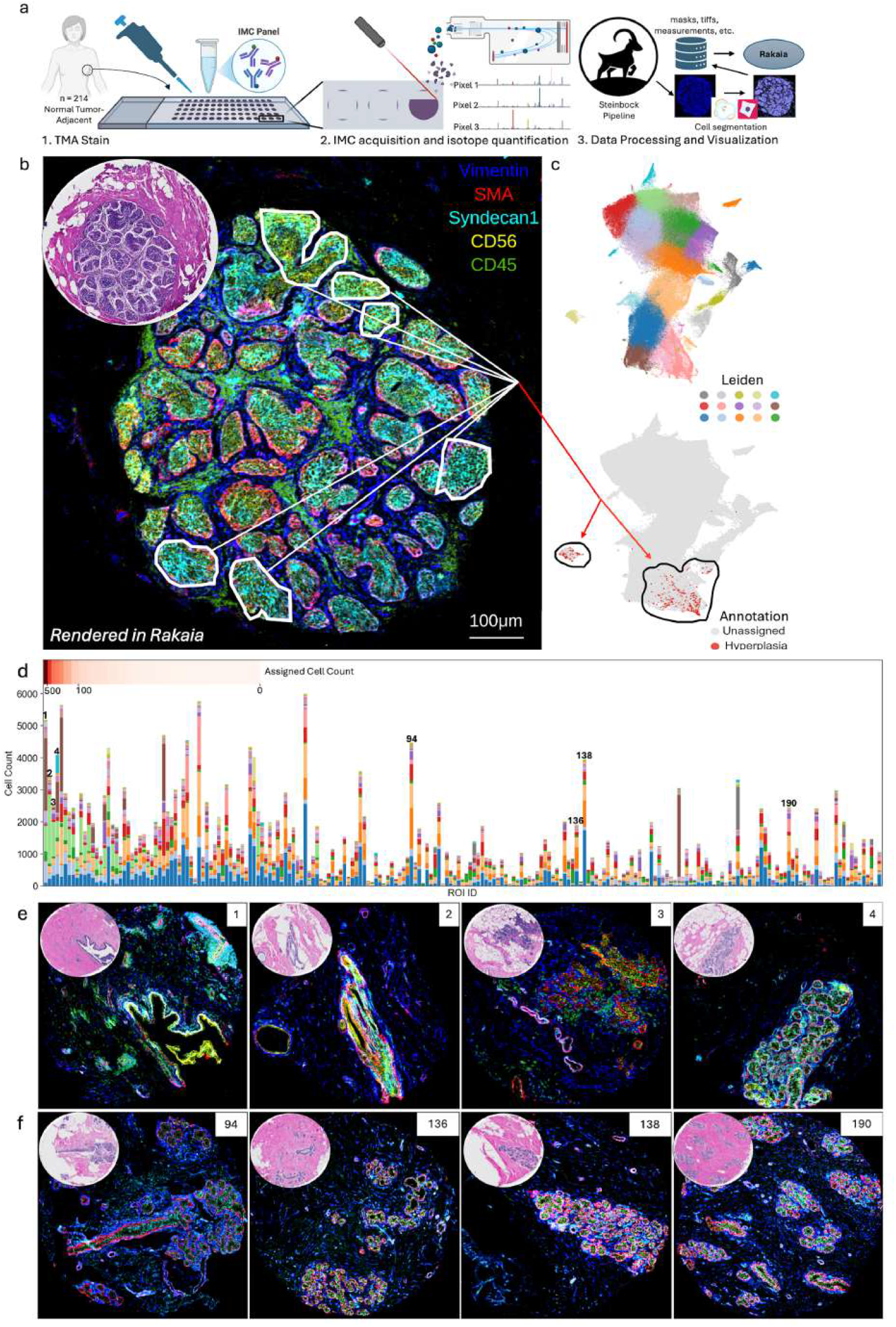
ROI prioritization in a multi-patient spatial cohort. **(A)** Schematic of imaging mass cytometry (IMC) workflow and data processing. **(B)** Manual annotation of sub-regions of tumour-adjacent breast tissue captures cells of interest, labeling them as “Hyperplasia” based on the query image. **(C)** UMAPs display Leiden clustering and the distribution of predicted cells classified in Rakaia, with assigned cells belonging to multiple clusters. **(D)** Total cell count for each Leiden cluster in non-annotated cores, ranked according to the number of cells predicted as hyperplastic. **(E)** Top four query regions from the Rakaia query gallery with the highest number of predicted hyperplastic cells. **(F)** Four regions with no predicted cells.

### Fast, interpretable, publication-quality images

Rakaia can render full-resolution, high-quality images using a canvas-style graph component. Users may browse a gallery of individual channel thumbnails, combining a selection of markers to create multi-channel blends for export or normalized visualization across entire datasets. Rakaia leverages various levels of on-demand data loading and transformation that makes rendering multiple large regions or whole slide images feasible using standard computational infrastructure (**Extended Data Fig. 2**). By modifying parameters for channel colour selection, pixel intensity scaling, and noise reduction, users can generate finely-edited images that are representative of experimental or biological variation. Scaling and normalization can be optimized for each image independently or across the dataset, enabling users to compare standardized images with greater interpretability. Rakaia provides multiple formats for image export enabling external dissemination, and cloud database or local storage of user selections, facilitating session reproducibility.

### Flexible annotations with matched object quantification

The main canvas provides tools to annotate individual pixels, segmented objects, or tissue structures with editing modes to zoom, pan, or draw enclosed freeform shapes to create annotations. These can capture groups of segmented objects or pixels to be integrated into quantitative visualizations with mapped object identifiers (**Fig. 2B**) and additionally be paired with interactive plotting to enable dataset-wide queries for objects with similar metrics. As an example, interactive selections on cell segmentation clusters in the Rakaia UMAP plot can be rendered across multiple images to reveal the spatial localization of cells with similar expression (**Extended Data Fig. 3**). Users can also visually gate objects directly in images by manually adjusting thresholds for the union or intersection of quantitative variables and interactively optimize the set of associated objects(**Extended Data Fig. 4**). Diverse user options for annotations with matched quantification profiles enable flexible phenotype-focused analysis to identify tissue architecture features or cellular niches (**Extended Data Fig. 4**), and can be combined with classification models for prediction across the entire dataset (**Fig. 2**). The combination of user-guided annotations and quantitative predictive modelling facilitates efficient, prioritized browsing of multi-image cohorts where assessing every region could be prohibitively time-consuming (**Fig. 2**).

To demonstrate Rakaia’s discovery workflow, we used pathology-guided tissue morphology to explore and guide cohort-level analysis of tumour-adjacent, non-malignant human breast tissue in a 43-dimension IMC experiment profiling >200 cores from >100 patients (**Fig. 2**). Visual image inspection identified one pathologist-confirmed example of abnormal breast tissue exhibiting in situ carcinoma and/or neoplasia (**Fig. 2B)**. Hand-drawn bounded annotations captured the cells within this hyperplastic tissue of interest, and a random forest classifier trained on the annotated image was applied to the remainder of the cohort, returning ranked images ordered on predicted cell content similarity (**Fig. 2B-D**). Visualized in Rakaia’s image gallery (**Fig. 2E**), top ranked query results show cores with the highest number of hyperplastic-like cells; including a core containing a ductal lesion ranked first, along with cores containing other pathological features such as microcysts and varying degrees of perivascular inflammation. Images absent for these cells rank towards the bottom of the query results and lack any notable pathological features (**Fig. 2F**). Thus, unified annotation, object quantification, and image ranking/prioritization in Rakaia enables efficient interrogation of a large data cohort, in this case guided by pathology defined tissue morphologies.

Rakaia facilitates in-depth exploration of increasingly scaled spatial datasets, adaptable to future data structures, emerging technologies, and novel analysis methods. Supporting both exploratory and focused analysis of diverse spatial data types, Rakaia is among the first platforms capable of converting complex, multimodal measurements spanning hundreds of images into coordinated, standardized visualizations with searchable elements and dynamic querying. Such a framework for comparative spatial analysis is necessary as experimentation continues to capture increasingly large numbers of features collected over vast image sets. With Rakaia, diverse groups of users will be able to realize new frontiers in the translation of high-dimensional multiplexed data into actionable biological insights, representing a new cornerstone to motivate high-throughput, multimodal research.

## Acknowledgments

This research was undertaken, in part, thanks to funding from the New Frontiers in Research Fund (NFRFE-2021-00278; KRC), Canada Research Chairs Program (HWJ: Tier 2 CRC-950-232945/2024 00231), NSERC Alliance (ALLRP 570709-2021 and ALLRP 598293-24; KRC and HWJ), NSERC Discovery (DGECR-2021-00366; HWJ), and OICR New Investigator (OICR IA-1-020; HWJ). KRC acknowledges support from the Canada Research Chairs program.

The biobanking of biological material was made possible through a collaboration with the Réseau de recherche sur le cancer (RRCancer) financially supported by the Fondation cancer du sein du Québec, and Oncopole, the FRQ cancer division, which receives funding from Merck Canada Inc., GSK, Pfizer and the Ministère de l’Économie, de l’Innovation et de l’Énergie du Québec. The RRCancer is affiliated to the Canadian Tumor Repository Network (CTRNet).

We respectfully acknowledge the Rakaia River of Aotearoa New Zealand, one of the multi-channel rivers of Te Waipounamu. This river holds deep cultural and genealogical importance to Ngāi Tahu, the tangata whenua of this region. We acknowledge their enduring relationship with the land and we commit to using this name with respect and care without seeking to define or appropriate its meaning. In doing so, we acknowledge our shared responsibility to support others by navigating access or easing resistance where it was previously difficult.

## Author Contributions

MW, KRC, and HWJ designed the project. MW and TTJ developed the software. GA and ELC designed, optimized, and performed IMC. GA performed data analysis. JLG provided user feedback for software optimization. CD provided patient samples. HKB provided pathology review. MW, GA, KRC, and HWJ generated figures and wrote the manuscript. All authors reviewed and approved the manuscript.

## Methods

### Software overview

Rakaia is a Python-based, web-deployed application that enables visualization and interactive analysis of multiplexed imaging and spatial transcriptomics datasets. It is available as a command-line interface (CLI) executable installable through Python package managers such as conda, pip or uv, as a containerized version using Docker, or as a binary distribution compiled for every major operating system (Windows, MacOS, Linux) using PyInstaller. These installation options enable Rakaia to be deployed locally, on premises, or cloud-based, and support access for a single or multiple users concurrently.

### Implementation/UI

Rakaia implements a micro web server using Flask v2.25 with Dash Bootstrap 1.4.1 web components providing the majority of frontend user interactivity through defined user input components; WSI rendering is provided through the openseadragon v5.0 (osd) Javascript module. Rakaia uses a common Python scientific computing stack of pandas v2.3.1 [29], numpy v1.26.3 [30], and numexpr 2.8.8 [31] on the backend server to perform array-based operations such as rasterization, scaling, filtering, and blending. Additional libraries in the Dash and Plotly software suite provide image and data visualizations components such as an image blend canvas, heatmap, and UMAP plot. The standard library vips [32] and the pyvips [33] Python binding are used to convert WSIs into deep zoom images (.dzi) for rendering in the browser with osd.

### Import/Configuration

Rakaia supports many commonly adopted multiplexed imaging and spatial technologies in their native data format (**Extended Data Fig. 5 and Table 1**). Data can be imported either from a local file path if the user has access to the server’s local file system, or through drag and drop file input widgets with any deployment configuration. Rakaia’s input components enable users to upload individual or multiple image files into a session provided that the files share a common marker panel; server side data transformations and lazy, on-demand file loading ensure that imaging data are loaded into memory only when requested for rendering by the user, enabling a standard laptop computer to support import and browsing of up to thousands of images in a single session.

### Upstream data processing

Rakaia directly supports outputs from several processing packages for spatial muti-omics, accepting common file formats for image normalization, segmentation, and quantification.

This includes raw or de-arrayed tissue microarray (TMA) images with matched segmentation masks from MCMICRO [33], spatial expression profiles in Anndata [35] format from sopa [36], and zarr directory stores for 10x Visium and Xenium assays generated through the spatialdata [37] framework. For IMC data in .mcd format, Rakaia supports the outputs from the steinbock [38] pipeline through either individual file uploads or in a single directory import command; we have developed a modified version of this pipeline made parallelizable by the Snakemake [39] workflow manager to process large IMC datasets across multi-core compute infrastructure. This modified pipeline performs IMC normalization/filtering, single cell/nuclei segmentation, and marker intensity quantification at single cell/nuclei resolution, with the addition of single-cell clustering using phenograph [40] and UMAP [41] dimension reduction and optimization of marker intensity scaling thresholds for image standardization during visualization. Rakaia also supports the ingestion of individual output files from other community-developed processing pipelines provided that they comply with the file formats described in Extended Data Table 1.

### Database for persistent user configuration

Users may optionally configure and connect to an Atlas mongoDB database instance using the pymongo [42] Python driver to store session configuration variables in the Document model format. Session configurations consist of user-selected blend parameters as well as browser session variables such as mask blending/opacity levels, legend/text settings, and custom channel aliases/labels. Saved configurations allow custom input values to persist across sessions and/or users, which can be shared through the database or in a JSON file. This decoupling of saving session parameters (which can often be summarized with just a few MB) separate from the raw images used to generate them (which in some extreme cases may be hundreds of GBs) makes image analysis highly reproducible while requiring minimal persistent storage space in a cloud database. Users can also deploy saved configurations across multiple datasets with the same biomarker panel, facilitating transfer of fine-tuned selections and enabling greater control for standardization of user selections across images belonging to large projects.

### Data loading/transformation

Rakaia uses a server side data transformation protocol backed by the Python pickle module to efficiently cache serialized large session data such as image layers for fast retrieval during callback invocation. This allows Rakaia to efficiently render and serve a multiplexed image to the user with minimal latency. Additionally, this enables ROI lazy loading, whereby raw imaging data for either a specific ROI or ROI channel (depending on the total pixel size of the region) are loaded into memory only when requested by the user; this scheme has the advantage of permitting users to import and parse hundreds of imaging regions in a single session while minimizing memory and CPU usage. This combination of lazy loading strategies and server side data transformation permits users to render selections from large imaging regions (e.g. ~ 50,000,000 total pixel count per channel with 400+ channels) (**Extended Data Fig. 2**) in the browser within a few seconds and toggle through a series of whole slide images with ease (**Extended Data Fig. 1**).

### Object expression classifiers

Rakaia includes object-level classifiers and clustering algorithms for summarized expression profiles stored in Anndata and CSV format. Classifiers (e.g. random forest from scikit-learn [43], and/or clustering (e.g. Leiden [44]) can be applied independently or combined iteratively to sets of quantified segmented objects matched across multiple ROIs, generating groups of object assignments that can be mapped back to images through the dataset gallery (**Extended Data Fig. 4**) or on the main canvas through category projection in a mask overlay (**Extended Data Fig. 3**). For classification modules, the input assumes a user has manually annotated at least one ROI of interest that serves as the object training set; the model then uses the training set to inform the classification on all other ROIs in the quantified set that were not part of the training split. Outputs from object classifiers and cluster algorithms are merged into the summarized expression profiles by object ID and may be overlaid in the UMAP plot (**Extended Data Fig. 4**) and/or exported in CSV format.

### Whole slide images (WSI) & alignment

Rakaia supports a variety of microscopy file types for rendering as WSIs (**Extended Data Table 1**); raw files converted into .dzi consist of an xml image manifest and sub-directories for individual tiles based on resolution level and are served through Flask static routing to the osd viewer when requested by the user. To enable users to perform guided exploration of WSIs relative to their multiplexed images, Rakaia supports pre-computed image alignment of WSIs (e.g. H&E stains) to spatial transcriptomics datasets such as 10x Visium and Xenium. For Visium NGS spot-based assays, WSI tissue positions in pixels are captured in the raw expression matrix, so direct mapping of Visium spot coordinates to H&E tissue regions is possible without any coordinate transformations (**Extended Data Fig. 2**). For in situ spatial transcriptomics assays such as 10x Xenium, Rakaia supports the use of precomputed affine matrices to perform coordinate alignment through matrix transformation. Users may zoom in on a region in their multiplexed image blend in the main canvas, and the WSI viewer will correspondingly update with the relative coordinate positions for the same sub-region in a matched WSI loaded with osd (**Extended Data Fig. 3**).

### Patient Samples

Human breast tissue was obtained from cancer-free, tumor-adjacent areas of resected samples and divided across 4 TMA slides, representative of 120 unique patients with two cores each. Accounting for core loss during the staining process, a total of 214 tissue cores were acquired and analyzed. Samples were stored in formalin fixed paraffin embedded (FFPE) blocks and two serial sections were utilized for hematoxylin and eosin staining in the first section and IMC staining in the second section. TMA construction and H&E staining [45] was performed by the Diorio Lab.

### Imaging Mass Cytometry Panel

A 43-marker, immune and epithelial antibody panel was prepared for staining across the 4 TMA slides. The list of antibody targets and clones are listed in **Extended Data Table 2**. Each antibody was conjugated to a lanthanide-series metal isotope of a unique mass to allow for detection via IMC. Conjugations were performed using the MaxPar antibody labeling kits (Standard Biotools) and according to the MaxPar labeling protocol.

### Imaging Mass Cytometry Staining and Acquisition

FFPE TMA slides were baked at 60C for 60 minutes. Tissue deparaffinization was performed in 3 × 10 min solutions of xylene and rehydrated in a graded series of reagent alcohol (reagent alcohol: deionized water 100:0 (x2), 96:4, 90:10, 80:20, 70:30, for 5 min each). Antigen retrieval was performed for 30 min at 90C in heat-induced epitope retrieval (HIER) buffer (10mM Tris Base, 1mM EDTA, pH of 9.2) using a Decloaking ChamberTM NxGen (Biocare Medical). Once cooled to room temperature, slides were incubated in blocking buffer (3% BSA and 5% horse serum in 0.1% TBST) for 1 hour. After blocking, slides were incubated in unconjugated primary antibodies diluted in blocking buffer overnight at 4C. Once rinsed in 3×5 min TBS solution, slides were incubated in metal conjugated secondary (anti-rat and anti-mouse) antibodies for 1 hour at room temperature. Rinsing was performed again in 3×5 min TBS solution to remove unbound secondary conjugates, after which slides were incubated in a cocktail of conjugated primary antibodies diluted in blocking buffer overnight at 4C. Next, slides were stained using iridium (a DNA intercalator, at 500 hM [JG1] for 5 min) and washed in 3×5 min TBS solution and then dried using pressurized air to prepare for IMC acquisition. Data was generated using an Hyperion XTi imaging system (Standard BioTools), laser-ablated in a 1um rastered pattern at 800 Hz and preprocessed using the commercial acquisition software (Standard BioTools).

### Imaging Mass Cytometry Data Processing

Raw IMC .mcd files were processed with through the steinbock pipeline v0.16.3 with the following parameters: cell and nuclei segmentation was performed using Mesmer from deepcell v0.12.6, and a hot pixel filter (hpf) of 50 was used; nuclear channels were set as Ir191 and Ir193, and the cytoplasm channel set as In115 and Yb174. The pipeline yielded single-cell and nuclei segmentation masks (.tiff) for all 214 ROIs, with segmented single-cell marker expression collected in .h5ad (Anndata) format. To perform normalization prior to clustering and UMAP, the segmented single cell expression data was censored at the 99.9th percentile to remove outliers and scaled by dividing each channel by the channel’s maximum value. No further transformations were performed. Cell annotation and plot generation were done using the Scanpy package, with unsupervised Leiden clustering performed to generate cellular clusters.

