## Supplementary Figures and Extended Data Tables for "Rakaia: interactive discovery of spatial biology at scale"

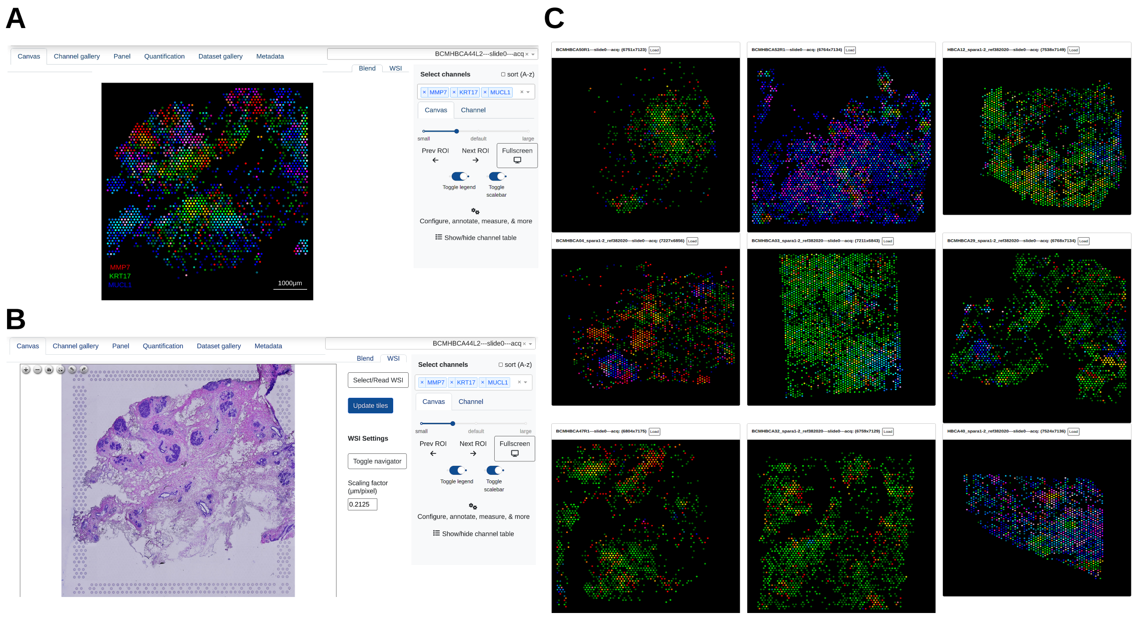


**Extended Data Figure 1 | Rakaia supports visualization of multiple spatial transcriptomics datasets with paired whole slide images. (A)** 10x Visium slides from 10 patients are simultaneously rendered with a subset of epithelial markers (MMP7, KRT17, and MUCL1) identified through spatial mapping and sub-clustering of unbiased spatial transcriptomics from the Human Breast Cell Atlas [46] **(B)** Matched H&Es may also be visualized alongside each spatial region, allowing the gene finding results of the study to be reproduced and confirming the histopathology associated with both luminal secretory (MMP7+) and luminal hormonal (MUCL1+) regions. **(C)** Rendering all of the spatial regions in the dataset gallery allows the user to prioritize subsequent images for main canvas focus or to visually interpret broad epithelial expression across the entire cohort. The multi-dataset rendering in Rakaia is extensible up to thousands of regions across all supported technologies.

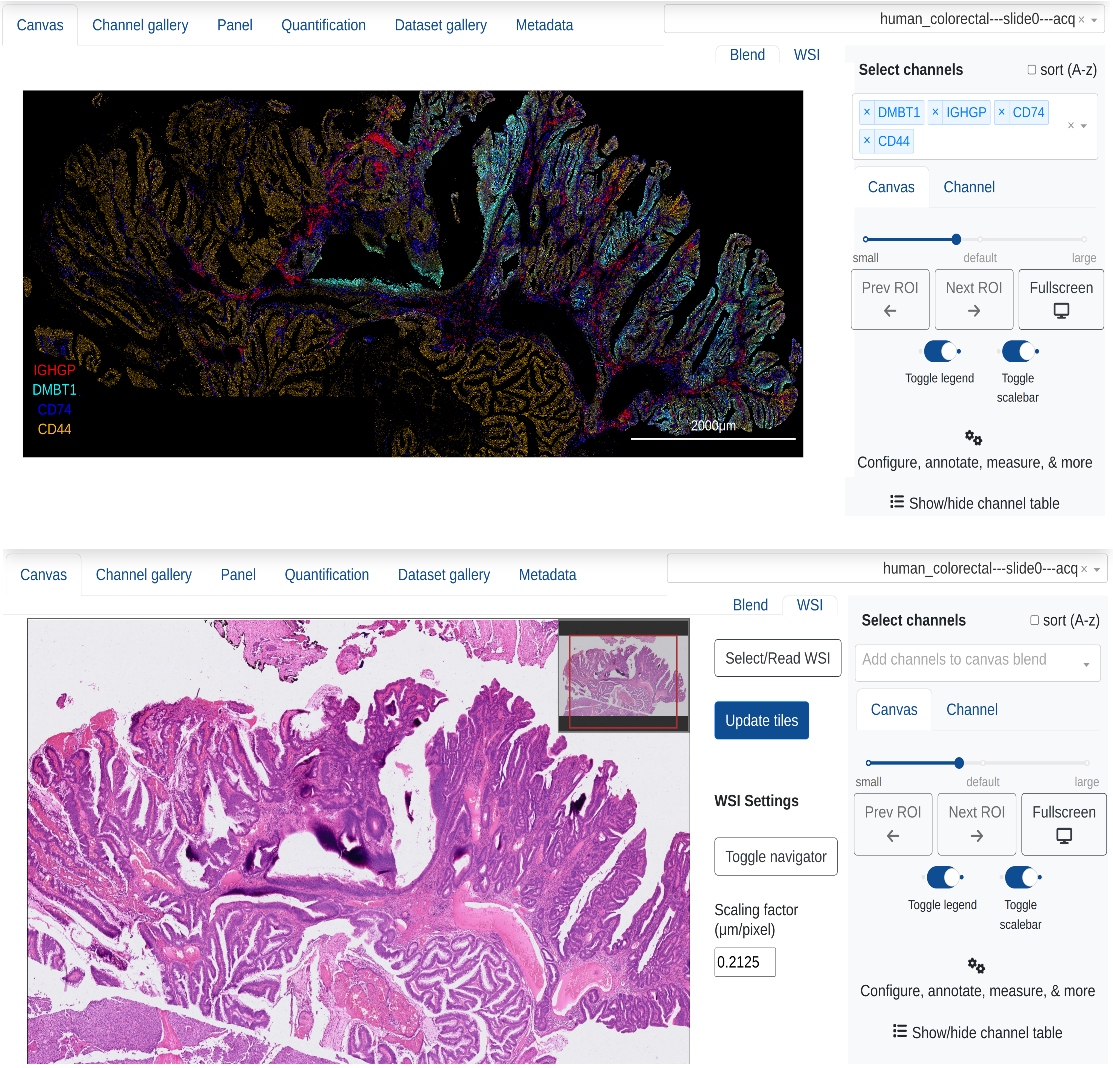


**Extended Data Figure 2 | Example visualization of a 10x Xenium in situ hybridization dataset with a >1cm dimension.** Rakaia can render large spatially resolved regions, such as human colorectal cancer (~0.4$cm^2$) profiled using the Xenium 380-gene immuno-oncology profiling panel from 10x Genomics. Differential expression combined with K-means clustering reveals markers for major immune and non-immune lineages in the tumour microenvironment, and a matched H&E WSI provides additional histopathology context for any transcript-based discoveries. With the use of pre-computed affine transformation matrices, zooming in on regions in the transcript image will apply a comparable zoom level into the corresponding subregion in the WSI. The dataset shown is not associated with any particular study, but highlights the flexibility of Rakaia to resolve spatial regions across imaging technologies and chemistries, augmented by histological imaging. URL: https://www.10xgenomics.com/datasets/ffpe-human-colorectal-cancer-data-with-human-immuno-oncology-profiling-panel-and-custom-add-on-1-standard

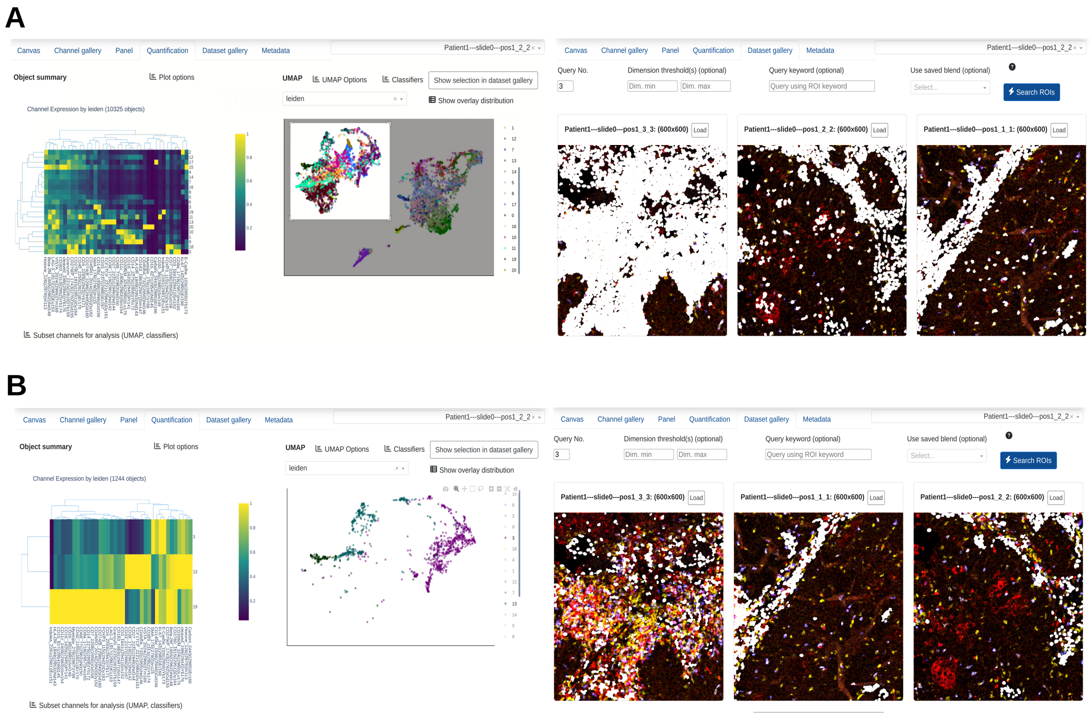


**Extended Data Figure 3 | Object expression profiles can be mapped to pan-ROI spatial locations interactively.** Tabular data structures for summarized object expression profiles can be imported and rendered in the quantification tab, supported by interactive plots such as heatmap and UMAP. **(A)** Zoom-in selections for sub-clusters in the UMAP permit users to map objects of shared UMAP dimensions across multiple ROIs in the dataset gallery, where query objects are shown in white overlaid on user-generated blends. Normalized object expression in the heatmap is also updated to the current user selection. **(B)** Similarly, categorical UMAP overlays can be subset to multiple values through interactive selection in the plot legend, providing users with further granularity for subsetting object groupings for mapping. Data shown are 3 squamous cell carcinoma of head and neck (SCCHN) ROIs from a single patient as part of the IMMUcan project [38].

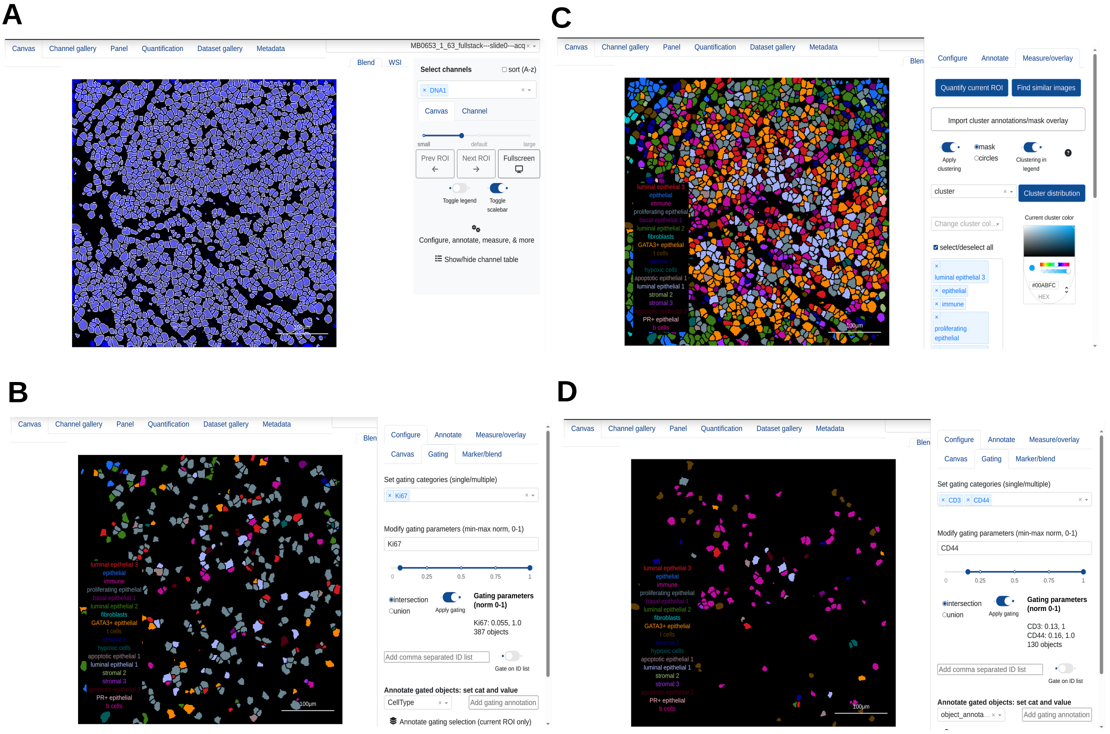

**Extended Data Figure 4 | Rakaia enables segmentation mask overlay, cell annotation projection and image-driven object gating. (A)** A representative ROI from the METABRIC breast cancer cohort profiled using IMC [47] is visualized with the cell segmentation mask overlaid on the nuclear (DNA) channel. **(B)** Cell-type assignments are projected into the overlaid segmentation mask, revealing the spatial context for cell phenotype combinations that may be indicative of cell-cell interactions **(C)** Visual-spatial gating for cells based on user-defined manual thresholds of Ki67 expression reveals the spatial distribution of predominantly proliferating epithelial cells. **(D)** Visual-spatial gating for cells based on the intersection of user-defined manual thresholds for CD3 and CD44 expression assists in identifying the distribution of broad immune and T cell-annotated populations.

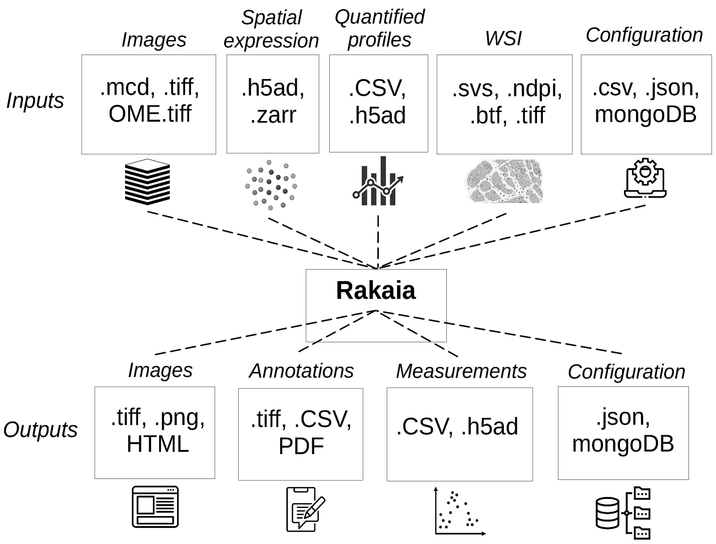


**Extended Data Figure 5 | Overview of Rakaia inputs and outputs**. Rakaia enables individual or single-command imports of raw images, spatial expression values, summarized object quantification profiles, and various session configurations from a variety of supported file types. Outputs from the application include standard file extensions for blended images, annotations in either image or tabular format that can be combined with matrix-held object measurements, and configuration files of the same format as the input. Optional user connections to a mongoDB database allow configurations to be saved to the cloud.

| File type extension(s) | Associated assay/data type | Associated Python package/module |
|---|---|---|
| .mcd | Imaging mass cytometry | readimc |
| .tiff, OME.tiff | Fluorescence, spectroscopy, immunohistochemistry, WSI, segmentation masks (multiple technologies) | tifffile, pyvips |
| .h5ad (Anndata) | Spatial transcriptomics (10x Visium, Visium HD, Xenium), expression profiles | scanpy, scipy, anndata, sopa |
| .svs, .btf, .ndpi | WSI | pyvips |
| Spatialdata/.zarr (directory) | Multiple technologies | spatialdata, spatialdata-io |
| CSV | Expression profiles, metadata, configuration | pandas |
| JSON | Metadata, configuration | json |

**Extended Data Table 1.** Rakaia can support multiple file types for spatially resolved data in native format. Various adopted file types for spatially resolved biology are supported in Rakaia, providing an interface for analysis of diverse spatial technologies and assay types. Associated Python packages supporting data parsing for on-demand data loading for images or spatial transcriptomic profiles.

| Antibody Target | Antibody Clone | Conjugated Metal |
|---|---|---|
| Vimentin | EPR3776 | Y89 |
| E-Cadherin | 36/E-Cadherin | In113 |
| Pan Cytokeratin | AE1, AE3, C11 | In115 |
| Kappa Light Chain | SP148 | La139 |
| HLA-DR | TAL 1B5 | Pr141 |
| CD303 | DLEC / CLEC4C / BDCA-2 | Nd142 |
| PD-L1 | CAL10 | Nd143 |
| CD28 | EPR22076 | Nd144 |
| CD15 | HI98 | Nd145 |
| CD45RA | HI100 | Nd146 |
| CD66b | G10F5 | Sm147 |
| ICOS | D1K2T | Nd148 |
| CD20 | L26 | Sm149 |
| CD68 | KP1 | Nd150 |
| CD4 | EPR6855 | Eu151 |
| CD8a | C8/144B | Sm152 |
| CD127 | EPR23747-333 | Eu153 |
| CD11c | EP1347Y | Sm154 |
| CD141 | EPR4051 | Gd155 |
| FOXP3 | 221D | Gd156 |
| TREM2 | D8I4C | Gd157 |
| GATA3 | L50-823 | Gd158 |
| TBET | E4I2K | Tb159 |
| FAP | AF3715 | Gd160 |
| Perforin | B-D48 | Dy161 |
| CD45RO | UCHL1 | Dy162 |
| Anti-rat | A18873 | Dy163 |
| Granzyme B | D6E9W | Dy164 |
| CTLA4 | CAL49 | Ho165 |
| Syndecan 1 | EPR6454 | Er166 |
| iNOS | SP126 | Er167 |
| Arginase | EPR6672(B) | Er168 |
| Anti-mouse | A28174 | Tm169 |
| TCF1 | C63D9 | Er170 |
| CD208 | EPR24265-8 | Yb171 |
| CD14 | SP192 | Yb172 |
| CD56 | EPR2566 | Yb173 |
| CD45 | D9M8I | Yb174 |
| CD1c | 3G1B3 | Lu175 |
| PD-1 | D4W2J | Yb176 |
| PNAd | MECA-79 | Pt195 |
| CD31 | RM1006 | Pt196 |
| SMA | 1A4 | Bi209 |
| CD3 | CD3-12 | Unconjugated |
| Gamma Delta TCR | H-41 | Unconjugated |

**Extended Data Table 2.** IMC antibody panel used for TMA staining. A complete list of all antibody targets used for the IMC panel along with corresponding clone information and conjugated metal isotope.
